# Parallel processing of natural images by overlapping retinal neuronal ensembles

**DOI:** 10.1101/2021.02.22.432289

**Authors:** Jesús Pérez-Ortega, Joaquín Araya, Cristobal Ibaceta, Rubén Herzog, María-José Escobar, Fernando Peña-Ortega, Luis Carrillo-Reid, Adrian G. Palacios

**Affiliations:** Centro Interdisciplinario de Neurociencia de Valparaíso, Facultad de Ciencias, Universidad de Valparaíso, 2340000, Chile; Department of Biological Sciences, Columbia University, New York, 10027, USA; Departamento de Electrónica, Universidad Técnica Federico Santa María, Valparaíso, 2340000, Chile; Instituto de Neurobiología, Universidad Nacional Autónoma de México, Juriquilla, Querétaro, 76230, México

**Keywords:** Retinal Neural Coding, Neuronal Ensembles, Visual System

## Abstract

Even though the retinal microcircuit organization has been described in detail at the single-cell level, little is known about how groups of retinal cells’ coordinated activity encode and process parallel information representing the spatial and temporal structure of changing environmental conditions. To describe the population dynamics of retinal neuronal ensembles, we used microelectrode array recordings that describe hundreds of retinal ganglion cells’ simultaneous activity in response to a short movie captured in the natural environment where our subject develops their visual behaviors. The vectorization of population activity allowed the identification of retinal neuronal ensembles that synchronize to specific segments of natural stimuli. These synchronous retinal neuronal ensembles were reliably activated by the same stimuli at different trials, indicating a robust population response of retinal microcircuits. The generation of asynchronous events required integrating a physiologically meaningful time window larger than 80 ms, demonstrating that retinal neuronal ensembles’ time integration filters non-structured visual information. Interestingly, individual neurons could be part of several ensembles indicating that parallel circuits could encode environmental conditions changes. We conclude that parallel neuronal ensembles could represent the functional unit of retinal computations and propose that the further study of retinal neuronal ensembles could reveal emergent properties of retinal circuits that individual cells’ activity cannot explain.

## Introduction

The retina is probably one of the most studied central nervous system structures since, technically, it is possible to preserve the whole structure *in vitro* and study its properties. The retina, located at the back of the eye, constitutes the first stage for the visual world’s neural codification. A photon flux integration occurred through a well-organized multilayer neural network (Wassle, 2004;Gollisch and Meister, 2010) supported by a variety of retinal ganglion cells (RGCs), which can be classified according to its genetics profile (Sumbul et al., 2014), morphology (Sanes and Masland, 2015), and physiology (Farrow and Masland, 2011). The physiological and collective behavior of RGCs (Nirenberg et al., 2001), e.g., neural synchrony (Shannon and Weaver, 1949;Mastronarde, 1983a;b;Levine, 1998;Barlow, 2001), has been proposed to encode different features of the visual world such as gratings (Hochstein and Shapley, 1976), sensitivity to the movement (Hammond and Smith, 1982;Frost and Nakayama, 1983;Born and Tootell, 1992), approaching (Munch et al., 2009) and differential movement (Olveczky et al., 2003). However, a still unresolved question is how the neural interaction between retinal neurons encode natural scenarios despite changing environmental conditions. Such problems cannot be solved considering the retina as an array of linearly connected neurons in a feedforward fashion. We investigated if the neuronal ensemble framework could provide insights into how different natural images are multiplexed and processed to reach a reliable neural output. Neuronal ensembles are groups of neurons with coordinated activity representing sensory stimuli, motor programs, memories, or cognitive states (Carrillo-Reid and Yuste, 2020a). It has been proposed that the activation of neuronal ensembles in the primary visual cortex could represent different features of natural images that signal relevant spatial and temporal information of the environment (Carrillo-Reid et al., 2015b). This way, several neuronal ensembles’ interaction could endow neuronal circuits with increased computational capabilities (Hebb, 1949;Yuste, 2015;Carrillo-Reid et al., 2016;Carrillo-Reid et al., 2019). Despite decades of research, the understanding of sensory systems is still missing definitive studies on the presence, structure, functionality, and dynamics of neuronal ensembles, in part because of Cajal’s and Sherrington’s postulates that affirmed that the individual neuron is the main computation module of neuronal circuits and that the flow of information was linear (Carrillo-Reid and Yuste, 2020b). The advent of multi-electrode array devices (MEA) (Meister et al., 1995) to record hundreds of cells simultaneously suggest that retinal population coding involves sophisticated and modular neural network architectures (Maffei et al., 1973;Movshon and Lennie, 1979;Smirnakis et al., 1997;Sharpee and Bialek, 2007;Gollisch and Meister, 2010;Greschner et al., 2011;Vidne et al., 2012;Trenholm et al., 2014). It has also been proposed using network models that retinal motifs connected in cascade with feedback connections could explain some emergent properties in retinal circuits (Real et al., 2017). These suggest that the retina’s information processing could be supported by neuronal ensembles recruited when appropriate visual stimuli are presented.

We performed population analysis on MEA recordings of the retina of degus, a diurnal dichromat visual rodent that has been used as a good model for the study of the visual system (Chavez et al., 2003;Palacios-Munoz et al., 2014;Escobar et al., 2018), to demonstrate that the synchronous activity of retinal cells define neuronal ensembles that represent diverse natural stimuli and that the retina’s functional organization allows the decomposition of visual stimuli information into parallel channels formed by functionally overlapping neuronal ensembles. Furthermore, the retina appears to be more than a passive counting photons device. It performs continuous processing of visual stimuli while adapting to the spatial and temporal properties of environmental changes.

## Results

### Retinal neuronal ensembles synchronize to natural stimuli

We recorded the activity of hundreds of RGCs simultaneously to investigate if the collective behavior of retinal neuronal ensembles can describe visual stimuli with different characteristics using MEA recordings of 256 channels. With this technique, the whole retina can be preserved and studied in vitro (Fig. 1A). To study retinal neuronal ensemble responses, we applied full-field scotopic or dark stimulus; photopic or light (cyan) stimulus; white noise (cyan and dark), and a natural movie (cyan) (60 Hz) corresponding to a 30 s sequence of a video recorded in a natural environment (Fig. 1B; see **Materials and Methods**). From the MEA recordings, individual RGCs were identified using spike sorting (Yger et al., 2018). A representative raster plot showing the population activity of individual RGCs demonstrated that the retina has different activation profiles to distinct visual stimuli (Fig. 1C). Firing rate was similar during scotopic and photopic conditions (0.9 ± 0.1 and 1.1 ± 0.1 spikes/s, respectively; mean ± SEM), but significantly increased during white noise and natural movie stimulation (6.2 ± 0.2 and 8.7 ± 0.2 spikes/s; mean ± SEM; Fig. 1D). However, the population’s coactivation profile also revealed synchrony peaks in response to natural stimuli (Fig. 1E). Natural movie evoked three times higher coactivations than white noise (0.6 ± 0.0 and 0.2 ± 0.0 z-score, respectively; mean ± SEM) even when the spike rate was only 1.4 times higher (Fig. 1D). These experiments suggest that the synchronized activation of RGCs groups could support the neural codification of relevant visual stimuli features (e.g., a natural movie). Specific retinal neuronal ensembles could encode different visual stimuli segments.

**Fig. 1.**
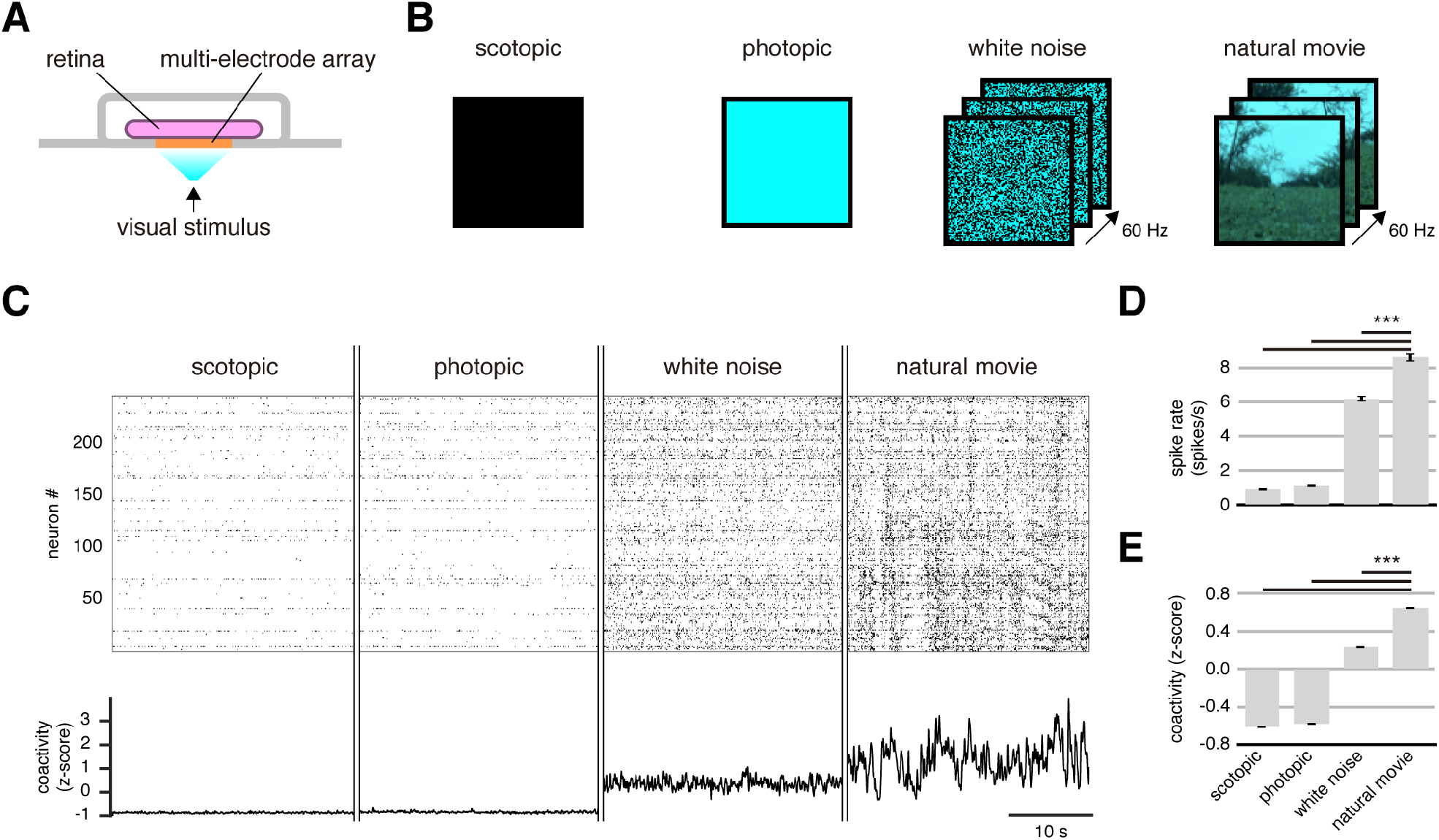
Natural stimuli evoke synchronized retinal neuronal ensembles. A) Schematic set-up for the MEA recording. B) Different stimuli used: scotopic, photopic, white noise, and natural movie. C) Representative raster plot for RGCs responses for different stimuli (Top) and the corresponding associated neural coactivity (Bottom). D) Spike rate for different stimuli: similar rates for scotopic and photopic stimuli (p = 0.78), but significantly higher during natural stimulation (p = 4×10^−9^). E) Average of normalized population coactivity for different stimuli. Note that the natural movie evoked three times higher coactivation than white noise for n = 867 neurons recorded from 4 different retinas; ANOVA test with post hoc Tukey-Kramer.

### Reliable activation of retinal neuronal ensembles

To investigate if retinal neuronal ensembles encode the same visual stimuli reliably at different times, we exposed the retina to the same movie (30 s length) 40 times. The raster plots reproduce similar RGCs activity patterns along with the trials and, consequently, similar coactivation profiles (Fig. 2A-C). To test reliability, we asked if it could be possible to predict what moment of the natural movie was presented by reading the neuronal responses. To do that, we trained a k-nearest neighbors (KNN) classifier with neural vectors from the first trial and classified the following trials. Neuronal vectors were created integrating the total number of spikes of each RGC at a given bin of time (Fig. 2D; See Materials and Methods). Prediction accuracy was dependent on the time window, above 75 ± 11 % (mean ± SEM) from a 500-ms bin (Fig. 2E, natural). Longer bins carry more data, so we asked whether the higher prediction was related to integrating the consecutive spikes during the time window. Then, we generated surrogated raster’s by shuffling 1-ms-bin neural vectors in time from the original raster (See Materials and Methods), but they were not reaching high prediction accuracy (Fig. 2E, surrogate). Thus, consecutive neuronal coactivation is needed to decode visual stimulation reliably. Moreover, the coactivation patterns evoked by the natural movie were also similar between different degus (0.56 ± 0.02 correlation, mean ± SEM; Figure 2F-G; for mice see Supplementary Fig. 1). We concluded that the same visual information is encoded by retinal neuronal ensembles reliably and robustly during different trials and evoked similar coactivation patterns even between different subjects suggesting hardwired ensemble connectivity.

**Fig. 2.**
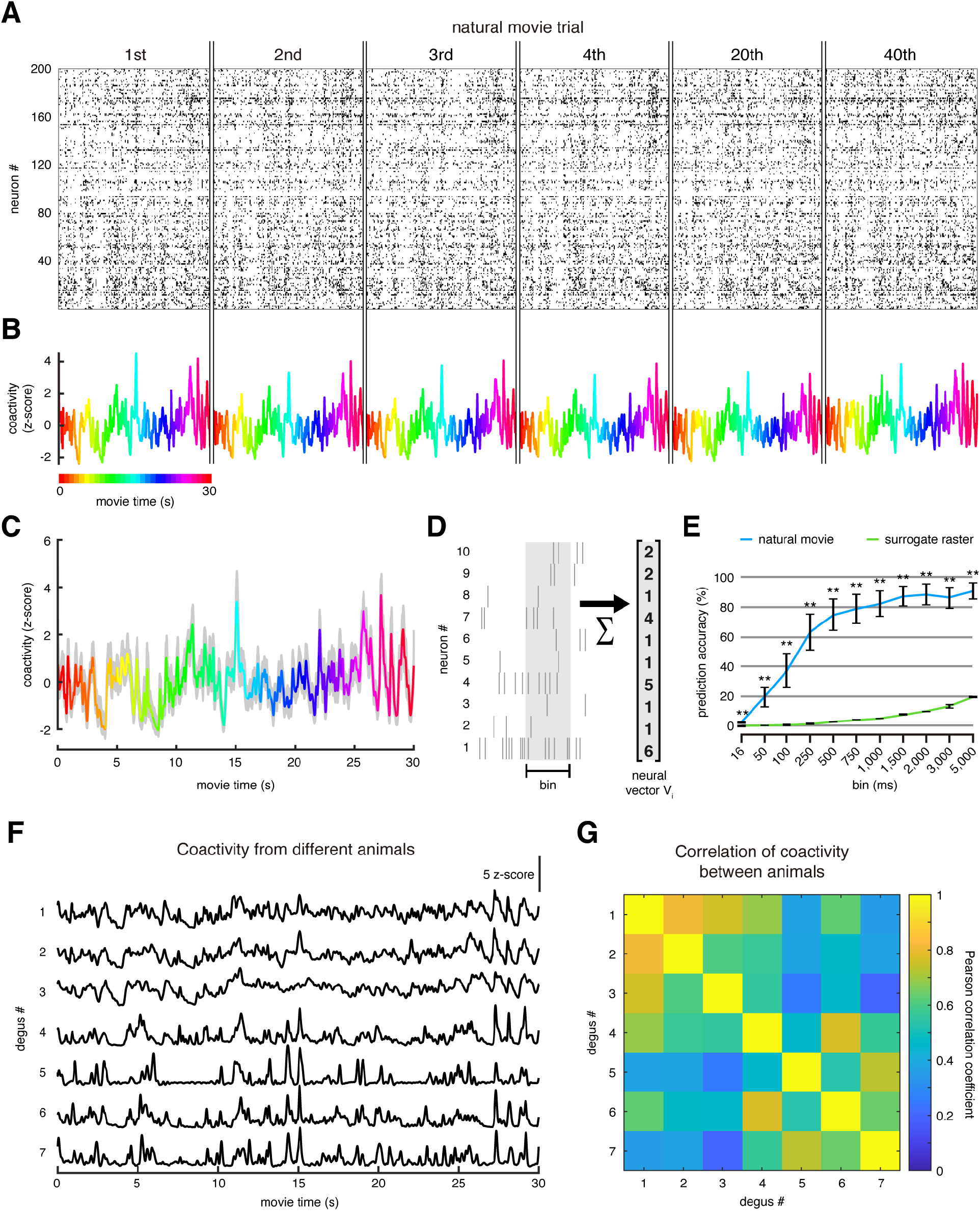
Retinal neuronal ensembles reliably respond to the same natural movie. A) representative raster plot for the 1st, 2nd, 3rd, 4th, 20th, and 40th trial (30-sec length). B) neural coactivity of RGCs and the trials in A; colors represent every second of the movie. C) coactivation of 40 trials overlapped (gray) and the average of all trials (colors). D) Schematic of how to extract neural vectors: we sum each neuron’s spikes during a given time bin. E) Prediction accuracy of the natural movie raster (blue) was significantly higher than its shuffled version (green) along with different time bins (16-ms bin p = 0.02; bins from 50 to 5,000 ms, p = 0.004). Mann-Whitney test; *p < 0.05, **p < 0.01. F) a Trial average of retinal coactivity evoked by the same natural movie recorded from different subjects (n=7). G) Pearson correlation coefficient between a trial average of retinal coactivity between different subjects (data on F).

### Retinal neuronal ensembles integrate time to encode natural stimuli

So far, we have demonstrated that the synchronous activity of retinal neuronal ensembles encodes relevant information of natural images. However, the time dependency of the responses and how fast such relevant information can be extracted remains unknown. We studied retinal neuronal ensemble responses to the same natural movie altered in time to respond to this question. We presented our natural movie during 20 consecutive trials with its original order, then 20 consecutive trials but played backward, i.e., frames in inverted order. Finally, we shuffled all the frames of the original natural movie (Fig. 3A). We observed that the original natural movie and the time-inverted natural movie evoked retinal neuronal ensembles with peaks of synchronous activity (∼60 % last longer than 100 ms and peak more than two z-score; p = 1×10^−11^ and p = 1×10^−11^, respectively), whereas most coactivity peaks during the shuffled movie were significantly shorter in duration and amplitude (∼70 % last less than 100 ms and below two z-score; p = 1×10^−11^ and p = 1×10^−11^, respectively) (Fig. 3B-D). Thus, the consecutive images in the natural movie stimuli were responsible for the retinal ensemble reliability activation. The coactivity profiles from the same stimulus repetition (standard or played backward) were as precise as we observed before. Still, unexpectedly, the evoked retinal neural ensembles for specific natural movie segments played forward, and backward were significantly different (Fig. 3E). Between the same stimulus repetitions, the correlation coefficient was above 0.9, but between forward and backward stimulus, the correlation coefficient was below 0.4 (Fig. 3F). Besides, the decoding accuracy (as computed in Fig. 2E) of the natural movie played forward was significantly lower using the data evoked played backward and vice versa (Fig. 3G). Retinal ensembles encoded differently the same natural movie stimulus depending if it was played forward or backward, suggesting that neuronal ensemble activity could be dependent on a physiological meaningful time window of structured visual stimuli including a complex optical flux sequence, changing eye view perspectives, left-right or up-down movement, etc.

**Fig. 3.**
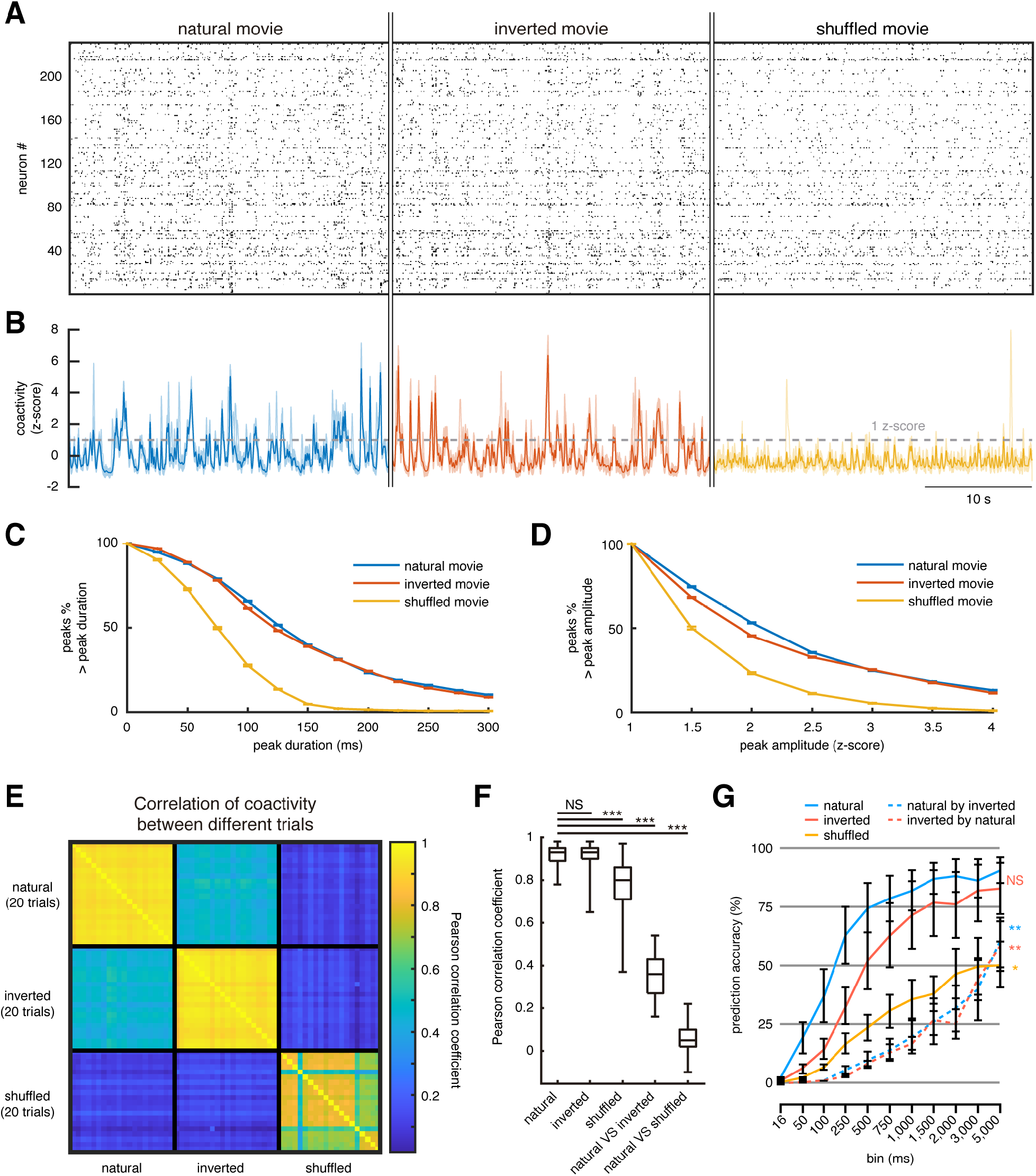
Retinal ensembles encode natural stimuli features by time integration. A) Example of a raster plot for a natural movie experiment with its frames in the original order (left panel), inverted order (middle panel), and shuffled order (right panel). B) Coactivity profiles overlapped from 20 trials (light colors) and its average (dark colors) during the natural movie (left), inverted movie (middle), and shuffled movie (right) condition. A dotted line at one z-score is placed to identify coactivity peaks for the following measures. C) Fraction of coactivity peaks lasting more than a given duration was significantly higher during natural or inverted than shuffled movie condition (p = 1×10^−11^ and p = 1×10^−11^, respectively). D) Fraction of coactivity peaks bigger than a given amplitude was significantly higher during natural or inverted than shuffled movie condition (p = 1×10^−11^ and p = 1×10^−11^, respectively). E) Pearson correlation coefficient between all trials of the example animal on B. F) Correlation average between trials of the same condition and between conditions. G) Prediction accuracy of the natural and inverted movie raster was significantly higher than shuffled movie raster along with different time bins (16-ms bin p = 0.02; bins from 50 to 5,000 ms, p = 0.004). Inverted movie prediction from natural movies or vice versa was significantly lower than the prediction using the same condition data. Mann-Whitney test; *p < 0.05, **p < 0.01; N.S. = Not Significant. Twenty trials/condition/subject (seven degus for a natural movie, and four degus for the inverted and shuffled movie).

On the other hand, coactivity evoked by the shuffled movie was also highly correlated between trials (> 0.75, Fig. 3F). Still, the decoding accuracy was significantly lower than the movie played forward or backward (Fig. 3G). In summary, our results suggest that structured visual spatial-temporal information (such as movements, accelerations, or stops during animal exploration of their natural environment) is critical to evoke the coactivity of retinal neural assemblies robustly. These results suggest the convenience of using a neuronal ensemble framework to study the response properties of retinal circuits.

### Retinal neuronal ensembles filter non-structured visual information

As we observed, the significant coactivation of the RGCs did not depend on the images per se but the natural sequence and structure of them. Then, we asked how many consecutive frames evoke a distinctive coactivity peak, i.e., retinal ensemble activation. We took 30 frames (500 ms) from the natural movie that evoked a coactivity peak, and we shuffled all of them (“shuffle”). We kept the first 1, 2, 3, 4, 5, 10 consecutive frames from the natural stimulus in the original order (Fig. 4A) and shuffled the rest without keeping them consecutively. We observed no significant response until five consecutive natural stimulus images were presented, producing a significant coactivity peak (Fig. 4B-C). We measured the area under the curve, in all conditions, where we observed the coactivity peak during a natural movie (200 ms bin, dotted line on Figure 2B). Coactivity peaks evoked by 5 and 10 consecutive images reached a similar area under the curve compared with population vectors evoked by natural stimuli (Fig. 4D). In our experimental conditions, five consecutive images from a natural stimulus were projected in 83.3 ms. Therefore, we concluded that this time is necessary to integrate and evoke a significant coactivity peak.

**Fig. 4.**
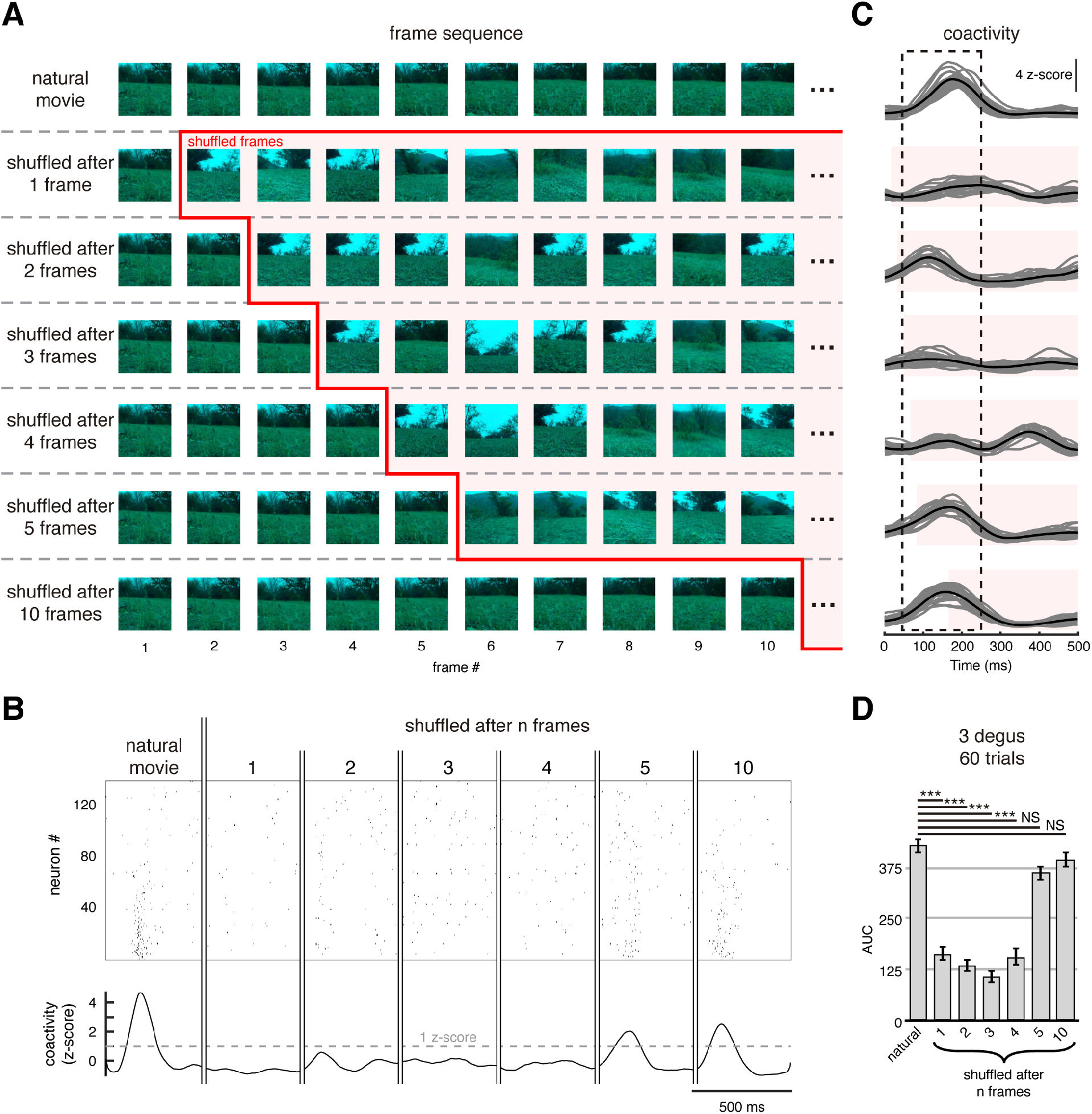
The minimal integration time window to evoke retinal neuronal ensemble activity. A) Different sequence of natural movie frames: original order (first row), which evoked a coactivation peak (see B and C); frame 1 in original order and the rest shuffled (second row); frames 1 and 2 in original order and the rest shuffled (third row); frames 1, 2, and 3 in original order and the rest shuffled (fourth row); and so on. B) Example of a raster (top) during a single trial of each sequence from A and its coactivity (bottom). C) Overlapped coactivity of 20 trials (gray) and its average (black) during the different sequence as in A. Shaded area is the time when the frames were presented shuffled. The dotted line represented when the coactivity peak was evoked during the natural movie’s original frame sequence. D) Area under the curve of the coactivity (dotted line on C) was significantly smaller during the first four shuffled sequences (p = 4×10^−8^). Still, it was not significant when we kept five consecutive frames (p = 0.6). Kruskal-Wallis test, with post hoc Tukey-Kramer: *** p < 0.001; N.S. = Not Significant. 20 trials/condition/subject (n = 3 retina).

### Spatially overlapping retinal neuronal ensembles encode natural stimuli information

To investigate the spatial distribution of retinal neuronal ensembles, we extracted the peaks of synchronous activity that signaled the same time segment of the natural movie (Fig. 5A) and compared the RGCs identity between different retinal neuronal ensembles (Fig. 5B). Then we created spatial maps identifying the location of RGCs that belong to different retinal neuronal ensembles (Fig. 5C). We observed a salt and pepper spatial distribution of neuronal ensembles and their receptive fields (Fig. 5D), suggesting parallel processing between neuronal ensembles’ activity. Retinal neuronal ensembles were heterogeneous with high overlap (Fig. 5E) and comprised mostly OFF RGCs (Fig. 5F). These experiments demonstrated that spatially overlapping retinal neuronal ensembles could multiplex different visual signal features to support a meaningful visual interpretation of the world. The activity profiles, observed at different trials, represent identified groups of RGCs defining retinal neuronal ensembles firing together, which activity depends on the visual contents’ structure.

**Fig.5.**
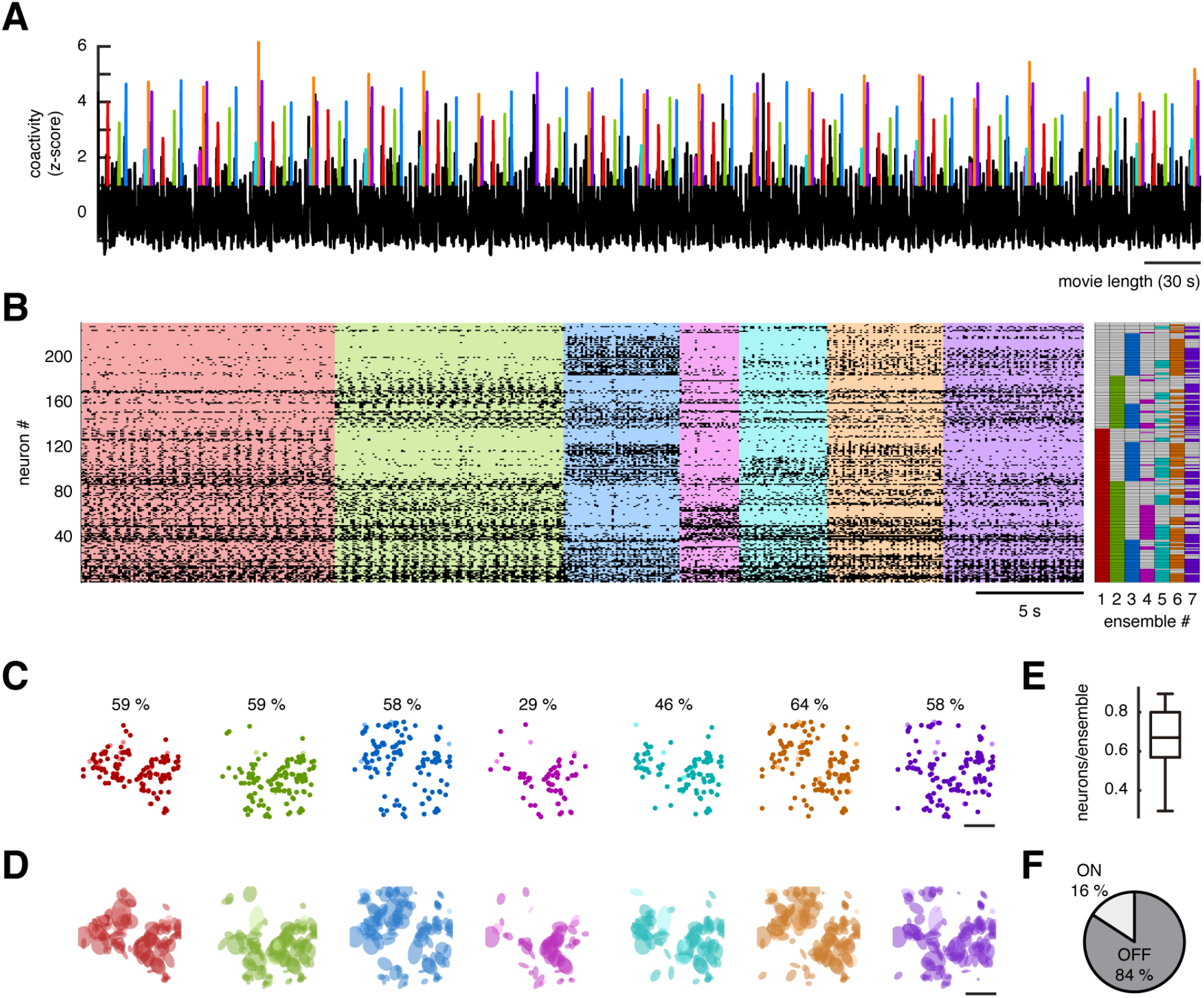
Overlapping retinal ensembles. A) z-scored coactivity of population vectors evoked by the repetition of natural visual stimuli 20 times. Each color denotes a specific retinal ensemble. B) Rater plot of sorted activity from A (Left) and identification matrix of neurons that belong to different retinal ensembles (Right). Note that retinal ensembles are heterogeneous and overlapping. C) Spatial maps of retinal ensembles highlighting the percentage of neurons they comprise. D) Receptive fields from RGCs. E) Fraction of the total neurons per retinal ensembles from C. F) Percentage of ON and OFF RGCs from retinal ensembles.

## Discussion

We used MEA recordings and population analysis to demonstrate that: (1) environmentally relevant visual stimuli recruits synchronous groups of cells that we defined as retinal neuronal ensembles, (2) the same retinal neuronal ensembles can be evoked by visual stimuli for several trials, showing the robust activation of ensembles, (3) the codification of spatial and temporal characteristics of visual stimuli requires the time integration of population activity that works as a filter for non-structured visual information, and (4) spatially overlapping, and flexible retinal neuronal ensembles represent changes in the visual environment in different animals.

### Visually evoked neuronal ensembles in the retina

It has been shown that the computational properties of the retina that allow the detection of directed motion rely on the precise connectivity between neurons (Briggman et al., 2011). Even though retinal neurons’ individual properties and connectivity patterns have been extensively studied, how motion computations are represented at the population level remains unknown. To explain the non-linear properties of retinal information processing, computational models formed by motifs connected in cascade with feedback connections have been implemented, demonstrating that retinal circuits’ emergent properties cannot be explained by single-cell feedforward connectivity (Real et al., 2017). Such evidence suggests that a change in the framework from a single cell to population analysis is mandatory to study the retinal circuits’ algorithms. Neuronal ensembles defined as groups of neurons with a coordinated activity that represent percepts, movements, or cognitive states (Carrillo-Reid and Yuste, 2020b) have recently proved useful to demonstrate the causal relationship between population activity and learned behaviors (Carrillo-Reid et al., 2019;Carrillo-Reid and Yuste, 2020a;Robinson et al., 2020), indicating that the ensemble framework could benefit the study of retinal neuronal microcircuits.

In this work, we demonstrated the use of population analysis to define retinal neuronal ensembles, providing evidence of collective and emergent neural activity at early stages of sensory processing, to study how groups of neurons maximize their responses to relevant spatial and temporal features related to the natural environment.

### Synchronous activity of retinal neuronal ensembles

We defined retinal neuronal ensembles constructing population vectors that capture the activity of all recorded neurons (Fig. 2). Such population vectors represent synchronous activity epochs that require the temporal integration of the visual information for at least 80 ms (Fig. 4). However, the mechanisms that allow the synchronization of retinal ensembles deserve further studies. As an initial step, we demonstrated that the blockade of gap junctions dramatically reduced the synchronization of the network activity (Supplemental Fig. 2A & 2B) and suppressed population responses’ ability to predict visual stimuli (Supplemental Fig. 2C) accurately. Thus, synchronization of retinal neuronal ensembles reconfigured by gap junctions could represent a mechanism to boost signal propagation throughout retinal circuits.

### Parallel processing by overlapping retinal neuronal ensembles

An unsolved question for retinal computation is how specialized retinal ensembles could be reliably activated despite visual stimuli’ initial state (Baden et al., 2018). One possibility is that different conditions recruit different ensembles, but the interaction between neurons that belong to different ensembles at intermediate levels convey the same meaningful output. Thus, for certain conditions, a significant output could be reached by structured spatial and temporal features captured by the interaction between retinal ensembles despite changes in the environment.

The fact that the same neurons could be part of different ensembles (Fig. 5) suggest that ensemble dynamics could reduce noise levels. On the other hand, the ensemble framework could provide insights on how parallel information is multiplexed in the retina. Cell participation in different ensembles and overlapping receptive fields demonstrates that functional retinal circuits are highly dynamic and reconfigurable to changing environmental conditions. Furthermore, shared members from different ensembles could activate overlapping ensembles in cascade, indicating that structured spatial and temporal visual information mat to robustly evoke an environmentally meaningful output, even between different subjects (Fig. 2F & 2G; Supplemental Fig. 1).

### The role of retinal neural ensembles in population coding

In the retina, the information is encoded both at the level of the single and population neurons. The complex circuitry given by two synaptic layers and more than 50 different cell types (Wassle, 2004;Sanes and Masland, 2015) generates networks with intrinsic variability due to noise, along with successive computations, internal network dynamics, and top-down chemical modulations, generating correlated activity response (Gardella et al., 2019). Population coding complements the information collected by single RGCs and imposes several advantages. It has been shown that correlated activity helps to increase the encoding accuracy. Noise correlated activity between excitatory and inhibitory inputs improves the time precision of retina response and its reliability (Cafaro and Rieke, 2010), and also, stimulus-dependent noise correlations enhance the accuracy of stimulus encoding (Franke et al., 2016;Zylberberg et al., 2016). Nevertheless, the results presented here suggest retinal neuronal ensembles obey input stimulus and are reconfigured accordingly. Moreover, this adaptation is not only to the input stimulus’s structure but also to the light intensity (Ruda et al., 2020). It remains a challenge to explore the variability of cells conforming to the neural ensembles depending on the input luminosity, establishing the conditions where the population processing dictates an advantage to encode the visual world.

## Materials and Methods

The experimental protocol for MEA recording we used here has been described before (Palacios-Munoz et al., 2014;Ravello et al., 2019). In brief, stimuli were generated using an appropriated code, where the timing of the images was controlled using a custom-built software based on Psychtoolbox for MATLAB.

### Animals

*Octodon degus* (n=7), both sexes, born in captivity at the Universidad de Valparaiso’s animal facility were maintained in light and temperature controlled with *ad-libitum* access to food and water. Dark-adapted animals, for 30 min, were deeply anesthetized with isoflurane and beheaded. Experimental procedures were approved by the Institutional Committee on Bioethics for Animal Research from the University (Certification # BEA050-15) and following the bioethics regulation of the Chilean National Agency for Research and Development (ANID). Degus are crepuscular/diurnal rodents with a high number of 30% of cones supporting a diurnal dichromatic vision (Chavez et al., 2003;Palacios-Munoz et al., 2014). To extend our observations, we also perform some experiments (n=3, 1 month old) on B6SJLF1/ mice as control.

### Electrophysiological recordings

Retinal ganglion cells (RGCs) responses were obtained using a Multi-Electrode Array device (Segev et al., 2004) (USB MEA256, with 100μm or 200μm electrode spacing / 30iR-ITO Multichannel Systems GmbH, Reutlingen, Germany) sampling at 20kHz. For recording, a small retina piece was mounted on a dialysis membrane array under perfusion with AMES medium bubbled with 95% 0_2_ + 5% C0_2_ at 33°C and the pH adjusted to 7.4.

### Spike sorting

Spike sorting analysis, to isolate a single RGC response from a multiunit recording trough several adjacent electrodes, was performed using SpyKING-CIRCUS (Yger et al., 2018). The algorithm considers for spike sorting were applied using spike-circus software the raw data passed through a high pass filter of 300Hz, to detects spike waveforms to identify and isolate cells from the full register. In this study, all cells that passed a signal/noise criterion and interspike interval violation of ≤ 2.5% were selected.

### Visual Stimulus

For stimulus display, a DLP projector was used using custom optics to reduce and focus an image onto the photoreceptor layer, projecting from the RGC side. A pixel’s size onto the photoreceptor layer was 4.7*µm* and an average irradiance of 70*nW/mm*^2^ (Newport 1918-R power meter, Newport Corporation, CA, USA. The DLP LEDs’ spectral emission has a peak emission at 460 nm and 520 nm (as measured with a USB4000 Fiber Optic Spectrometer, Ocean Optics), both in the range of covers the dichromatic, green, and the U.V. cones, vision of degus (Chavez et al., 2003). A natural movie that reproduces the natural environment of degus was obtained using a small telecommand four wheels device, home built, with a mounted camera (GoPro HD Hero 1, 1/2.3” sensor, 16:9 @ 60 fps, 1280 x 720p) allowing pre-programmed displacement with smooth scanning up, down, left, right, slow, fast movements to emulate natural displacement of degus exploration in nature (RA Vasquez, 2002).

### Obtaining coactivity

We use timestamps of unitary recordings at 20 kHz from MEA to build a binary matrix R of size N x S, where N is the number of neurons and S the number of samples, representing time. In the R matrix, one represents if a neuron has at least one spike and 0 if not in a given time. The bin resolution was 1 ms. The number of active neurons in each sample were counted (sum of columns of R matrix), obtaining the coactivity vector (1 x S) or coactivity signal. The coactivity was smoothed with a moving average filter of 100 ms to detect coactivity peaks easily. We should indicate we got no time-shift in the filtered signal.

### Defining neuronal ensembles

We considered significant coactivity peaks that are above a threshold (P < 0.01) of the signal. The threshold is determined by generating a surrogate shuffled raster (Carrillo-Reid et al., 2015a;Yger et al., 2018). Time-shift shuffles were used, where all neurons preserve their firing pattern but each one with a different temporal shifting. We generated 1,000 shuffled raster plots and their cumulative distribution function (CDF) of their coactivity. The number of co-active neurons at the value of 0.99 of the CDF was taken as the threshold. The null hypothesis is that the activity of neurons is independent, and then the coactivity occurs randomly. Spikes in a given significant coactivity peak were counted for each neuron and converted to a neural vector V of size N x 1 (Sasaki et al., 2007). To identify if the same activation of neurons produced similar neural vectors, the Euclidean similarity matrix S of size P x P were computed for all vectors (*V*_1_, *V*_2_, …, *V*_*P*_):

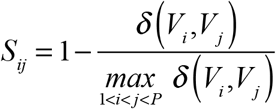

where P is the number of vectors and delta is the Euclidean distance. A similarity matrix was then used in a hierarchical cluster analysis using Ward linkage (this kind of linkage had the best faithfulness of the whole ensemble and a cascade of neighboring other consistency in our hierarchical trees). We used the contrast function described in (Beggs and Plenz, 2004).

### Shuffling raster

We shuffled the raster’s activity of each animal during the first movie trial maintaining the activity pattern of each 1-ms bin. Then, we generated the following surrogated raster trials keeping the same shuffling order. Doing this, we tested if the prediction accuracy was based on the consecutive activity patterns.

### Shuffling frames of a natural movie

To test if the sequence of frames in the natural movie were responsible for evoking significant coactivity peaks, we stimulated the retina with the same frames. Still, we shuffled the apparition order every 5 seconds. In shuffle 1, all frames were shuffled with a resolution of 1 frame. In shuffle 2, all frames were shuffled with a one frame resolution, except the first two consecutive frames. In shuffle 3, all frames were shuffled with a resolution of 1 frame, except the first three consecutive frames, and so on with shuffle 4, 5, and 10.

## Acknowledgments

This work was supported by grants from ICM-Chilean Science Millennium Institute ANID grant P09-022-F and FONDECYT # 1200880 to AGP. AFOSR Grant FA9550-19-1-0002 (MJE and AGP).

## SUPPLEMENTARY INFORMATION

**Supplemental Figure 1.**
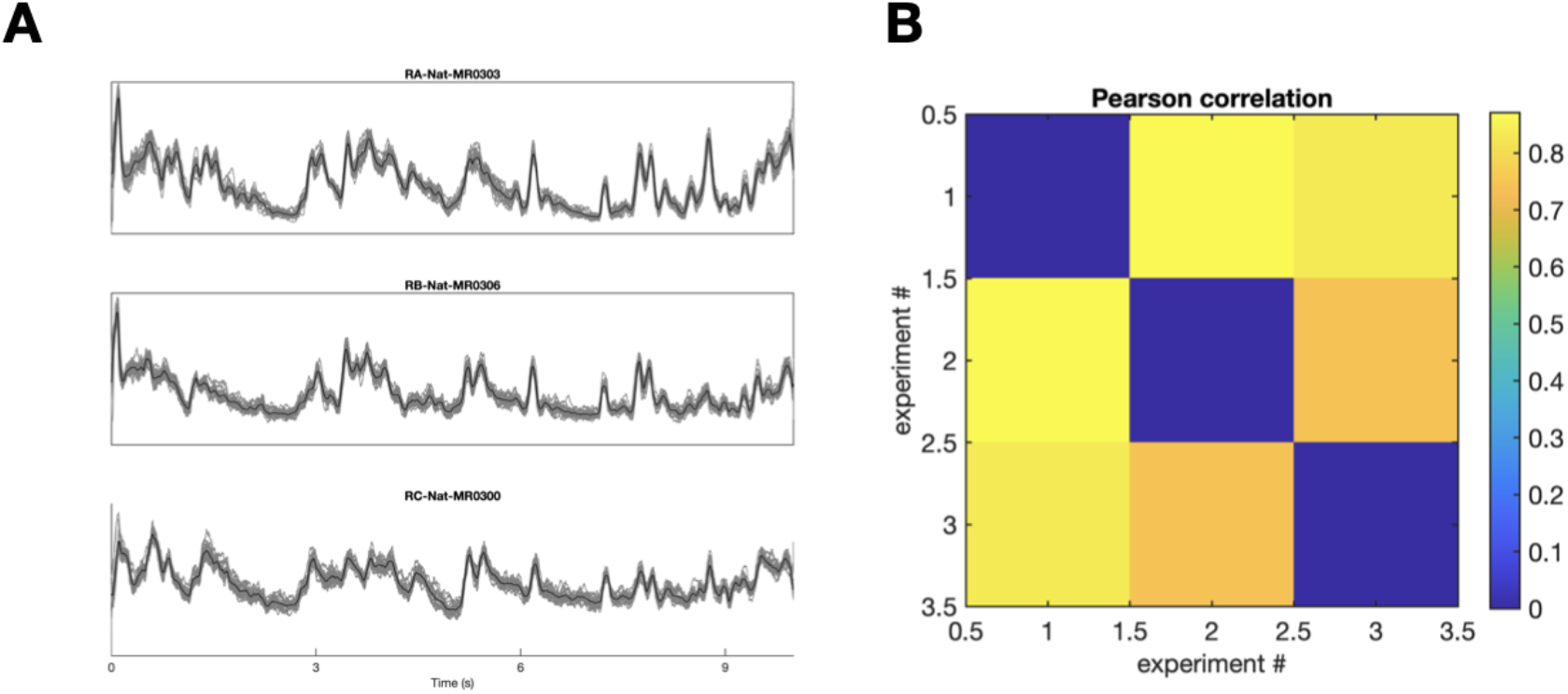
Retinal responses from different animals. A) Average population response to natural stimuli in different animals. B) Confusion matrix of Pearson’s correlation coefficient between different animals. Note that the structure of the responses is preserved in different animals.

**Supplemental Figure 2.**
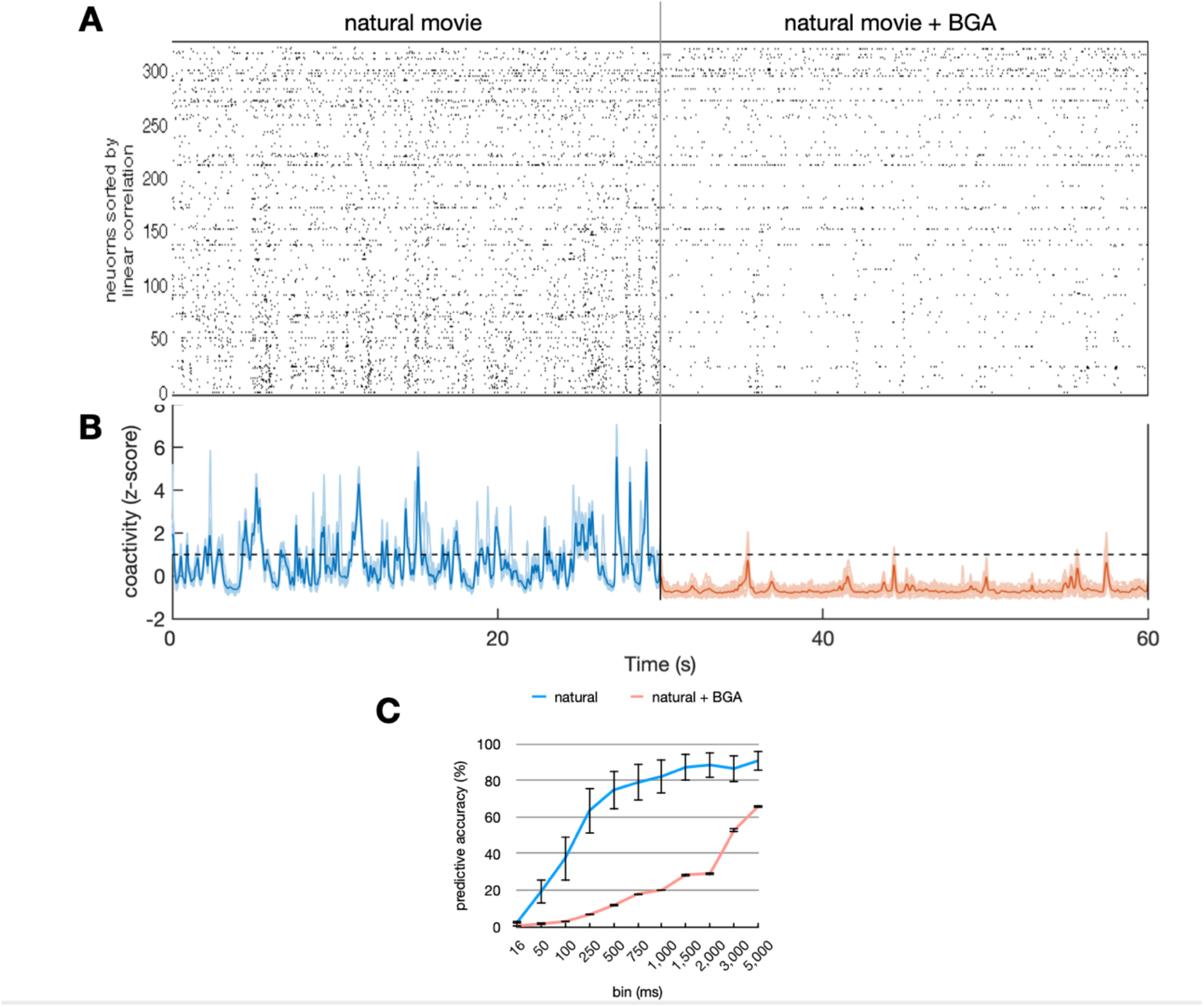
Role of gap junctions in retinal structured activity. A) Raster plot of retinal responses to natural stimuli before and after the blockade of gap junctions with BGA. B) z-scored coactivity in control and BGA experimental conditions. C) Predicting accuracy is impaired after the blockade of gap junctions.

